# Three-Dimensional Fluid-Structure Interaction Reveals Trade-offs Among Oxygen Delivery, Shear, and Washout in Pediatric Roller-Pump Cardiopulmonary Bypass

**DOI:** 10.64898/2026.10.26.761989

**Authors:** Yunpeng Tu, Yi-Ting Yeh, Alex de Lecea, Kevin Charette, Lyubomyr Bohuta, Vishal Nigam, Juan C. del Álamo, Christina L. Greene

## Abstract

**Objective:** To determine how roller speed, tubing occlusion, and hematocrit jointly regulate oxygen delivery, shear, and washout in pediatric roller-pump cardiopulmonary bypass.

**Methods:** A three-dimensional, closed-loop fluid-structure interaction model resolved roller-tube contact, tubing deformation, and blood flow. Forty simulations combined 4 roller speeds (38-95 revolutions per minute), 2 occlusion levels (77% and 83%), and 5 hematocrit levels (15%-50%). Cycle-averaged flow, oxygen delivery, peak shear stress, and viscosity-normalized relative residence time were quantified.

**Results:** Roller compression produced retrograde leakage through the occluded gap, localized peak shear stress exceeding 100 pascals, and adjacent stagnation and recirculation. Flow increased nearly linearly with roller speed and was higher at greater occlusion, with little hematocrit dependence. Higher speed and occlusion increased peak shear stress while reducing relative residence time; higher hematocrit further increased peak shear stress. Similar oxygen delivery could arise from substantially different flow environments: among 10 conditions providing oxygen delivery of approximately 9-12 milliliters per minute, peak shear stress ranged from approximately 25 to 70 pascals. Within conditions satisfying an illustrative minimum oxygen-delivery threshold, lower hematocrit permitted speed and occlusion settings yielding lower peak shear stress and relative residence time.

**Conclusions:** Pediatric roller-pump settings that provide similar oxygen delivery can expose blood to markedly different shear and washout environments. Joint evaluation of these metrics may guide selection of speed, occlusion, and hematocrit to meet a target oxygen delivery while reducing shear and improving washout.

## INTRODUCTION

Approximately 40,000 children receive open-heart surgery yearly in the US, requiring cardiopulmonary bypass (CPB) for circulatory and respiratory support^1^. Roller pump-based CPB remains the dominant method for pediatric cases due to its compatibility with small circuits and low priming volume^2^. Despite its indispensable role, CPB is associated with serious postoperative complications, including systemic inflammatory response syndrome and multi-organ dysfunction^3,4^. Multiple factors have been implicated in these adverse outcomes, including blood contact with artificial surfaces, temperature modulation, ischemia-reperfusion injury, transfusion exposure, and excessive mechanical stress on circulating cells^5–11^. Supraphysiological flow shear is increasingly recognized as a contributor to CPB- or mechanical circulatory support-associated morbidity via hemolysis, platelet activation, and pro-inflammatory leukocyte activation^12–17^. Recent mechanobiology studies showed that monocytes exposed to CPB-relevant shear upregulate inflammatory pathways^13,14,16,17^.

Current CPB operation focuses on maintaining adequate indexed oxygen delivery (DO_2_i) to prevent ischemic injury, such as acute kidney injury, while minimizing clinically observable complications. Yet reported safe DO_2_i thresholds vary across pediatric cohorts^18–22^, and target hematocrit (Hct) remains controversial: Jonas and colleagues identified a threshold near 23.5%^23–25^, whereas other studies support diverse strategies^26–29^. Moreover, roller-pump performance cannot be characterized by peak flow or oxygen delivery alone. In practice, oxygen transport is regulated by adjusting pump speed, Hct, and occlusion. These parameters, along with other circuit parameters such as tubing diameter, shape the stresses experienced by circulating cells. Thus, similar oxygen delivery indices may mask distinct, incompletely characterized biomechanical insults. From a mechanical perspective, roller-pump CPB is a high-occlusion form of peristaltic pumping, a classical transport mechanism in which a traveling compression wave displaces fluid through a deformable conduit^30^. The advancing wave generates transient and spatially localized tubing deformation, flow acceleration, and elevated shear stress^31–33^. It may also create disturbed transport regions characterized by recirculation and inefficient washout, exposing circulating blood cells to prolonged unphysiological flow in addition to short bouts of elevated shear^34,35^. Despite their potential biomechanical relevance, shear and transport-related metrics are rarely incorporated into perfusion decision-making.

Computational modeling offers a promising approach to investigating unphysiological flow stresses in CPB roller-pump circuits^31–33,36,37^. Existing studies follow two approaches: lumped-parameter models^36,37^, which capture circuit-level pressure and flow-rate waveforms but lack spatial resolution, and computational fluid dynamics (CFD) analyses^31–33^, which resolve local flow structures, but largely rely on two-dimensional domains with simplified models of tubing deformation. Neither approach captures the three-dimensional interaction among roller motion, tubing deformation, and flow. In addition, the combined effects of roller speed, occlusion, and hematocrit have not been systematically examined. Consequently, how these parameters jointly regulate oxygen delivery and flow-mediated insults remains poorly understood.

Three-dimensional fluid-structure interaction (FSI) analysis can feasibly resolve coupled roller-tube contact, deformation, and flow in roller pumps^38^. However, its potential for analyzing and optimizing CPB operation has not been realized yet. Here, we developed a three-dimensional FSI model of a pediatric roller-pump CPB circuit and conducted a systematic campaign of simulations varying the operating conditions. Our analysis identifies operating strategies that balance oxygen transport and biomechanical insults such as shear exposure, and disturbed washout.

## METHODS

### Roller pump computational model

Figure 1 situates the computational model within a representative CPB circuit (Fig. 1A) and shows the LivaNova S5 pump head on which it was based (Fig. 1B). The pump comprised a 150-mm-diameter raceway, two opposing 30.5-mm-diameter rollers with adjustable occlusion, outlet tubing clamps, and eight cylindrical guides (four per side). A 0.09-mm clearance between the guides and tubing outer wall prevented out-of-plane displacement. During each 360° rotation, every tubing section was compressed twice, once by each roller.

**Figure 1.**
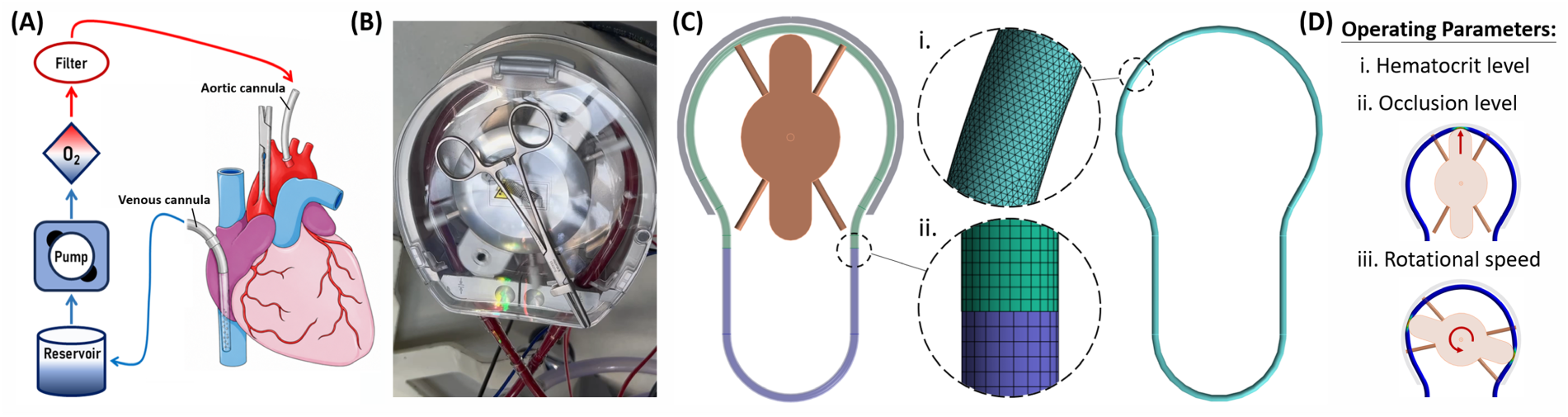
Roller pump cardiopulmonary bypass (RP-CPB) circuit, pump geometry, computational model construction, and investigated operating parameters. **(A)** Simplified schematic of the RP-CPB circuit. **(B)** LivaNova S5 Roller Pump used as the basis for model construction. **(C)** Computational model including solid-domain assembly (left) and closed-loop fluid domain (right). Insets show **(i)** the unstructured tetrahedral fluid mesh and **(ii)** the bonded interface between the deformable (green) and fixed (purple) tubing segments. **(D)** Investigated operating parameters: **(i)** hematocrit levels of 15%-50%, **(ii)** roller occlusion levels of 77% and 83%, and **(iii)** rotational speeds of 38-95 rpm.

The system was modeled in Ansys 2022 R2 using a solid domain for the pump assembly and tubing, and a fluid domain for the circulating fluid. The solid domain comprised the backplate (gray), dual-roller assembly and guides (brown), and tubing (green, blue; Fig. 1C). The tubing inner diameter matched the nominal clinical size (4.76 mm), whereas its outer diameter (6.35 mm) followed commercially available flexible PVC tubing. The tubing was divided into a deformable raceway segment (green) and a fixed segment (blue) connecting the downstream and upstream ends of the raceway to form a closed loop, while the fixed segment represented the tubing immobilized by the pump clamps. The two segments, of lengths 400 mm (green) and 263 mm (blue), were joined by bonded interfaces (Fig. 1C-ii). The fluid domain occupied the tubing lumen and formed the same closed loop, allowing roller-induced deformation to generate flow without prescribed inlet or outlet conditions.

### Fluid-structure interaction simulation framework

Roller-induced tubing deformation was simulated with the Transient Structural module in Ansys Mechanical APDL. To transition smoothly to steady operation, the rigid roller assembly was prescribed three successive motions: translation toward the tubing to establish the target occlusion (initial engagement, 0<t<0.4 s); a linear increase in angular speed from rest to the prescribed value (rotation ramp, 0.4<t<3.4 s); and constant-speed rotation thereafter (t>3.4 s; Supplementary Fig. 1).

The deformable tubing was modeled as an isotropic, linearly elastic material (Young’s modulus, 40 MPa; Poisson’s ratio, 0.48; density, 1500 kg·m^-3^)^39,40^. The tubing was discretized using the MultiZone method with quadratic 20-node hexahedra and quadratic wedges as required in curved regions. A nominal 0.8-mm element size yielded 98,067 nodes and 15,840 elements. One quadratic element was used through the wall because the analysis targeted global lumen deformation rather than through-thickness stresses. Frictionless contact was imposed at the tubing-roller, tubing-guide, and tubing-backplate interfaces.

One-way fluid-structure interaction (FSI) was implemented using ANSYS System Coupling through a surface-to-surface interface between the inner surface of the deformable tubing and the corresponding fluid-domain boundary. The fixed tubing segment was excluded from coupling. At each coupling step, tubing-wall nodal displacements were transferred from the structural solver to a System Coupling dynamic-mesh boundary in Fluent. A two-way FSI simulation confirmed that flow-induced pressure gradients negligibly affected tubing deformation, supporting the one-way approximation. The structural and fluid solvers were synchronized using identical time steps.

### Computational fluid dynamics model

Flow was simulated in ANSYS Fluent 2022 R2 using a transient, three-dimensional, finite-volume formulation under the assumptions of incompressible, Newtonian flow governed by the Navier-Stokes and continuity equations. A no-slip boundary condition was imposed at the tubing wall, whose motion was prescribed through the FSI coupling. The closed-loop tubing geometry enabled the flow to be solved without prescribed inlet or outlet boundary conditions. No turbulence model was used.

The fluid domain (Figure 1C, right) was discretized using an unstructured tetrahedral mesh with a nominal element size of 0.4 mm, yielding 304,542 nodes and 1,596,693 elements. Transferred wall motion was propagated using Spring/Laplace/Boundary Layer smoothing and implicit local cell- and face-based remeshing, with a residual criterion of 10^-5^, length scales of 0.24-0.59 mm, and maximum cell and face skewness values of 0.79 and 0.57, respectively. The mesh provided approximately 47 elements across the undeformed tubing and 12 across the maximally occluded section.

Convective terms were discretized with third-order MUSCL, using PRESTO! pressure interpolation and PISO pressure-velocity coupling; time integration was second-order implicit. Up to 30 nonlinear iterations were performed per time step, with convergence defined by scaled residuals below 10^-6^ for continuity and 10^-5^ for momentum. A 0.0004-s step was used during roller engagement and the rotation ramp (t<3.4 s) and during subsequent flow development until flow rate and domain-averaged and peak shear stress became periodic. This required 3, 4, 6, and 7 cycles at 38, 57, 76, and 95 rpm, respectively. The step was then reduced to 0.00025 s for an additional 1.4 pump cycles, and metrics were calculated over the final complete cycle. The Courant-Friedrichs-Lewy (CFL) number remained below 0.4 throughout the domain except for the occlusion region, where it averaged 2.2.

Mesh and time-step sensitivity were evaluated at 83% occlusion, 50% hematocrit, and 95 rpm. Nominal element sizes of 1.2, 1.0, 0.8, 0.6, 0.4, and 0.3 mm were tested. Time steps were scaled proportionally to element size to maintain comparable CFL numbers. Relative to the 0.3-mm reference, differences for the 1.2-, 1.0-, 0.8-, 0.6-, and 0.4-mm meshes were [8.71, 8.4, 7.17, 5.31, and 2.47] % in flow rate and [58.65, 45.61, 31.69, 24.88, and 2.91] % in peak shear stress. Simulations with dt = 0.0000625, 0.000125, 0.00025, 0.0005, and 0.001 s were then compared under the same condition. Relative to the reference simulation with dt = 0.0000625, differences for dt = 0.000125, 0.00025, 0.0005, and 0.001 s were [0.08, 3.37, 8.99, and 10.38] % in flow rate and [3.26, 3.52, 10.21, and 20.99] % in peak shear stress. Based on these results, the 0.4-mm mesh and 0.00025-s time step were selected for final simulations.

All simulations were performed on the Hyak high-performance computing cluster at the University of Washington.

### Operating parameter space

Roller occlusion was defined as

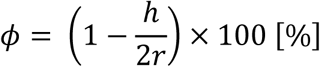

where ℎ is the minimum lumen gap prescribed during roller engagement and *r* is the undeformed tubing inner radius. Two occlusion levels were prescribed: ***ϕ****=* 77% and 83%, corresponding to ℎ ≈1.1 and 0.8 mm. Four constant roller speeds (*Ω* = 38, 57, 76, and 95 rpm) were investigated. For each pair of ***ϕ*** and *Ω* values, five hematocrit (Hct) levels ranging from 15% to 50% were considered. Blood density was fixed at 1060 kg·m⁻³ as its Hct dependence was negligible compared with that of viscosity.

Hematocrit-dependent viscosity (*μ* units: mPa·s) values were determined from Long et al.’s empirical regression^41^ for pediatric blood under an oscillatory strain of 5, with Hct expressed as a percentage: μ=2.58+4.211×10^-4^Hct^2.36^

### Post-processing

Flow visualization and data extraction were performed using Ansys CFD-Post 2022 R2, and Secondary analyses and associated graphs were performed with MATLAB R2022b.

Assuming total oxygen saturation, a hemoglobin concentration linearly proportional to Hct, and neglecting the small contribution from dissolved oxygen, oxygen delivery^19^ (DO_2_, units: mL·min^-1^) was determined from the cycle-averaged flow rate *<u>Q</u>* (L·min^-1^) and hematocrit as *DO*_2_ = 4.624 *<u>Q</u> Hct*.

Flow shear stress (*τ_s_*) was calculated as the product of the dynamic viscosity (*μ*) and the shear-rate (*γ̇*), 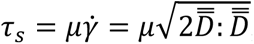, where *D̿* is the strain-rate tensor, defined as the symmetric part of the velocity gradient tensor. At each time point, the maxima of *γ̇* and *τ_s_* were identified within the fluid domain. Then, the cycle-averaged peak shear rate (*<u>S</u>_max_*) and stress (*<u>τ</u>_max_*) were calculated as the time average of these spatial maxima.

Relative residence time (RRT; units: Pa^-1^) is usually defined as

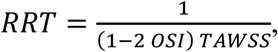

where *OSI* and *TAWSS* denote the oscillatory shear index and time-averaged wall shear stress^42,43^. Because blood viscosity varies with Hct in our study, we used the modified RRT (μRRT; units: s)

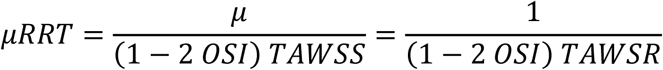

where *TAWSR* (units: s^-1^) is the time-averaged wall shear rate, *TAWSR* = *TAWSS*/*μ*. Elevated μRRT corresponds to near-wall regions experiencing recirculation and reduced washout. To characterize RRT variation along the raceway while reducing sensitivity to isolated extreme values, we reported the 75^th^ percentile of the μRRT at each position.

## RESULTS

### Roller compression generates high shear, backflow, and recirculation

To examine how CPB operating conditions may affect circulating cells, we analyzed tubing deformation and intraluminal flow across combinations of roller speed (*Ω*), occlusion (*ϕ*), and hematocrit (Hct). Figure 2 illustrates roller-induced deformation, localized flow shear, and altered blood transport at peak shear for the case with the highest occlusion, hematocrit, and roller speed (*ϕ* = 83%, Hct=50%, *Ω* = 95 rpm).

**Figure 2.**
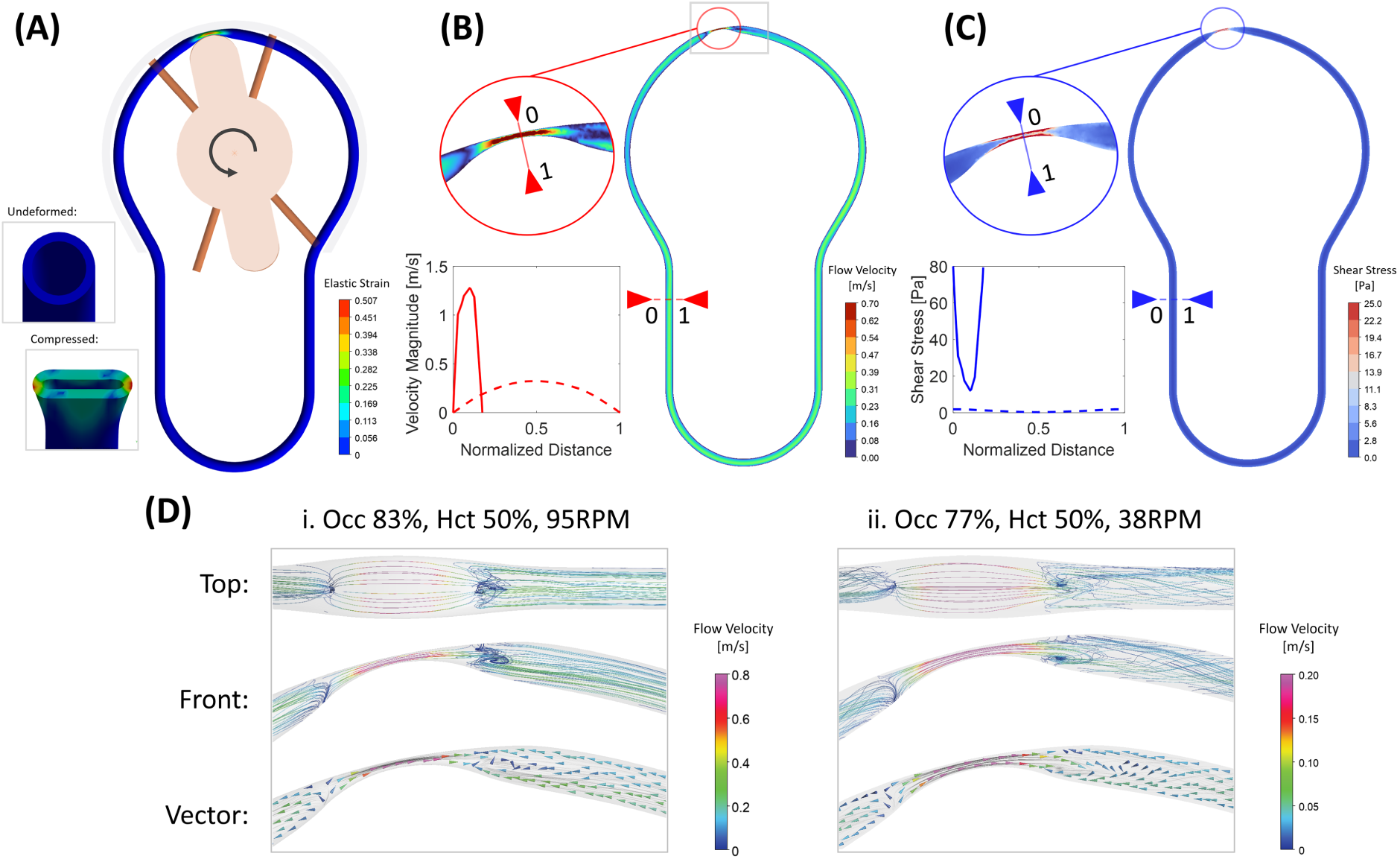
Roller compression generates localized high shear, backflow, and recirculation. Results of ϕ= 83%, Hct=50%, Ω= 95 rpm are shown at peak shear stress. **(A)** Equivalent elastic strain in the deforming tubing; insets show the undeformed and compressed tubing cross-sectional geometry. **(B, C)** Velocity and shear stress on the same cross-sectional plane, respectively; insets show each field within the occluded region and corresponding profiles across the compressed gap and free tubing. **(D)** Velocity-colored streamlines and vector maps near the occlusion for **(i)** ϕ=83%, Hct=50%, Ω=95 rpm and **(ii)** ϕ=77%, Hct=50%, Ω=38 rpm, illustrating how backward gap flow interacts with forward bulk transport to produce stagnation and recirculation.

Peak shear stress occurred when the roller compressed the tubing approximately 11° from the top of the raceway during counterclockwise rotation. At this timepoint, the tubing’s circular cross-section was bent and flattened into the obround shape as shown in Figure 2A inset, widening laterally from its nominal radius of r=3.175 mm to r=4.317 mm. The resulting three-dimensional deformation of the tubing wall reached maximum local elastic strain values of approximately 0.5. Moreover, because the tubing was not fixed to the backplate, the entire tubing displaced within the raceway, in line with operating-room observations (Supplementary Movie 1). These features are typically not captured by previous two-dimensional computational models of roller pumps^31–33^.

Figure 2B shows the cross-sectional velocity field and inset profiles across the occluded gap and free tubing, with transverse position normalized by the nominal tubing diameter. At 83% occlusion, the gap produced a sharply peaked profile with a maximum velocity of 1.545 m/s, approximately four times the free-tube value (0.334 m/s), where the profile was broader and symmetric. The free-tube velocity agreed with previous computational studies^31–33^, which is notable given that it emerged from the closed-loop pump-circuit dynamics rather than being prescribed via inflow/outflow boundary conditions. The Reynolds number was Re=266, which was below reported values (500-1400) consistent with our smaller pediatric circuit. Peak gap velocity was near the lower end of previous reports^31–33^, likely because the prior studies used greater normalized occlusion in larger tubing^32,33^ and two-dimensional domains that prevented lateral widening of the compressed lumen.

Localized acceleration through the occluded gap concentrated shear stress in the roller-compressed region (Figure 2C), where it exceeded 100 Pa versus approximately 2 Pa in most of the remaining tubing. These values agreed with some previous studies^32,33^ but were below those reported by Mulholland et al.^31^, consistent with the peak-velocity differences above.

Flow through the occluded gap reversed relative to the net flow direction (Figure 2D), while the flattened lumen produced lateral expansion and convergence of the instantaneous streamlines. This three-dimensional pattern created a stagnation region on the counterclockwise side and a recirculation region on the clockwise side of the occlusion, consistent with flow trapping and prolonged residence time. Reversed flow and trapping are characteristic features of high-occlusion peristaltic pumping^30^.

All simulations exhibited similar disturbed flow patterns, with only modest case-to-case variations. At high occlusion, hematocrit, and roller speed (*ϕ* = 83%, Hct=50%, *Ω* = 95 rpm; Figure 2Di), stronger net forward flow rapidly realigned streamlines on both sides of the roller, confining disturbances to a small region. At low occlusion and roller speed (*ϕ* = 77%, Hct=50%, *Ω* = 38 rpm; Figure 2Dii), weaker forward flow produced a larger recirculation region.

### Roller speed, occlusion, and hematocrit differentially regulate oxygen delivery and shear generation

We examined how roller speed, occlusion, and hematocrit jointly regulate oxygen delivery (DO₂), cycle-averaged flow rate (<u>*Q*</u>), peak shear rate (*<u>S</u>_max_*), and peak shear stress (*<u>τ</u>_max_*) across 40 simulations (Figure 3). The campaign included two occlusion levels (*ϕ* =77% and 83%), four roller speeds (*Ω* = 38, 57, 76, and 95 rpm), and five hematocrit levels (Hct=15-50%). Because roller speed is the primary parameter adjusted during CPB, results are plotted against speed, with each curve representing fixed hematocrit and occlusion.

**Figure 3.**
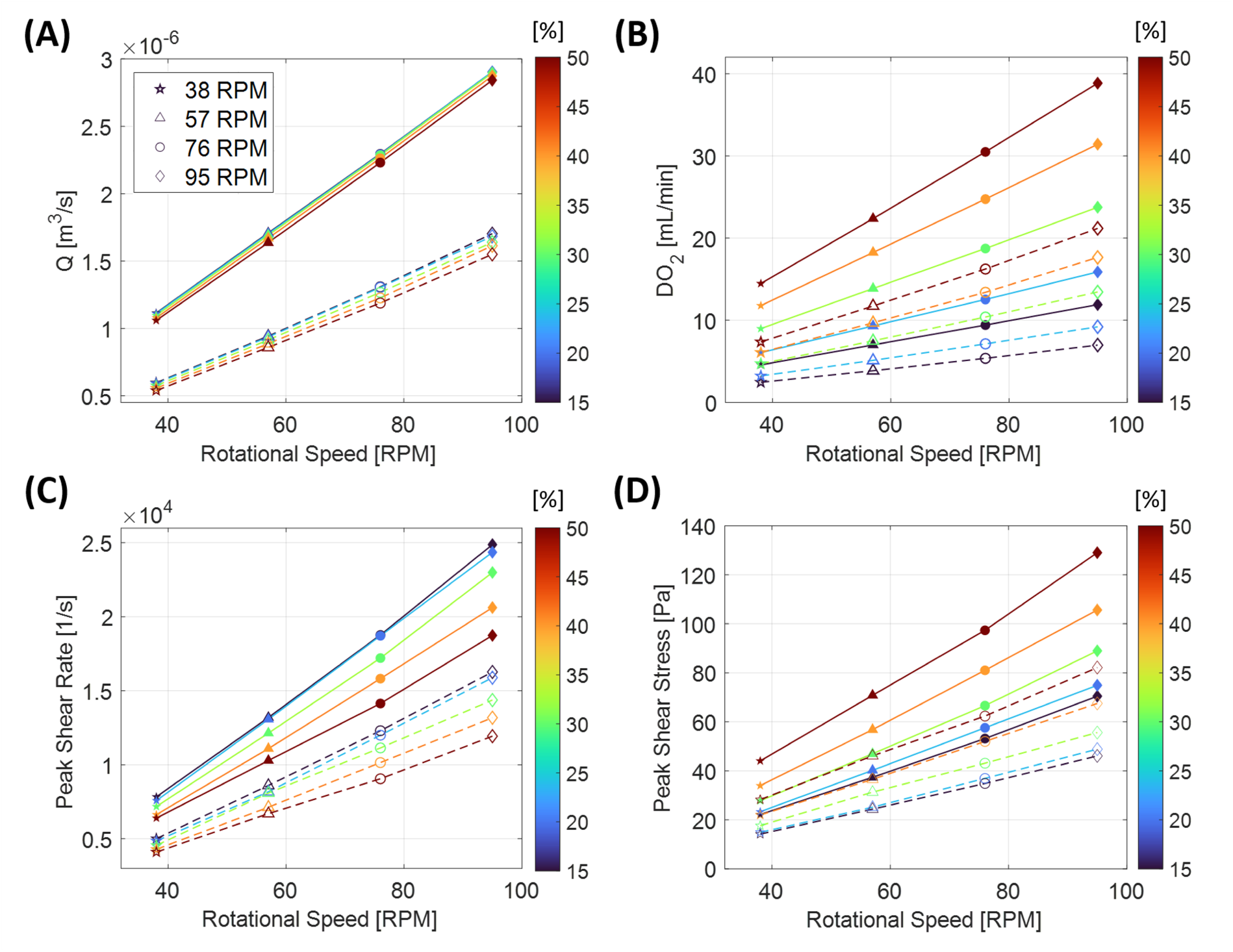
Effects of CPB operating conditions on cycle-averaged flow transport, oxygen delivery, and flow shear. **(A)** Cycle-averaged volumetric flow rate (*<u>Q</u>* [m^3^/s]), **(B)** oxygen delivery (DO_2_ [mL/min]), **(C)** peak shear rate [s^-1^], and **(D)** peak shear stress [Pa] as functions of roller rotational speed (*Ω*, rpm) under combinations of two occlusion levels (77% and 83%; open and filled symbols, respectively), four rotational speeds (38, 57, 76, and 95 rpm; symbol shapes), and hematocrit levels ranging from 15% to 50% (symbol color). Trend lines connect cases with fixed hematocrit and occlusion to illustrate the dependence of each metric on *Ω* (dashed and solid lines represent 77% and 83% occlusion, respectively).

Figure 3A shows that *<u>Q</u>* increased almost linearly with *Ω* over the investigated rpm range. This behavior is consistent with classical low-Reynolds-number peristaltic pumping theory, which predicts *<u>Q</u>*∼*r*^2^*R_p_Ωf*(*ϕ*), where *r* is the inner tubing radius, *R_p_* the pump radius, and the occlusion factor *f*(*ϕ*) increases from 0 to 1 as *ϕ* increases from 0 to 100%. The theory also includes a negative correction associated with viscously dominated backflow through the occlusion^35^. As *ϕ* approaches complete occlusion, this correction vanishes, making *<u>Q</u>* independent of fluid viscosity. Accordingly, increasing *ϕ* in our simulations increased the slope of the *<u>Q</u>*-rpm relationship, whereas hematocrit exerted only a minor influence. The weak dependence of *<u>Q</u>* on hematocrit suggests that the operating conditions were in the high-occlusion regime, where viscous backflow is small. The predicted flow rates were also consistent with previous simulations^31–33^.

Hematocrit was a significant determinant of oxygen delivery (Figure 3B), producing greater separation among the DO₂ vs. rpm curves than among the flow-rate curves. Importantly, overlap among these curves showed that comparable oxygen delivery could be achieved through different parameter combinations; for example, 20% hematocrit at *ϕ* = 83% produced nearly the same DO₂ vs. rpm relationship as 40% hematocrit at *ϕ* = 77%. Thus, alternative operating strategies could achieve the same oxygen-delivery target while generating different shear environments, potentially allowing biomechanical insult to be minimized.

To determine whether these alternative operating strategies produced different shear environments, we examined peak shear rate and stress. *<u>S</u>_max_* increased with roller speed and occlusion (Figure 3C), reflecting faster backflow and steeper velocity gradients, respectively. It varied little across lower Hct values but tended to decrease at higher Hct because increased viscosity resisted backflow through the occlusion. A similar dependence on *Ω* and *ϕ* was observed for *<u>τ</u>_max_* (Figure 3D), but its hematocrit dependence was reversed. Because shear stress scales with the product of viscosity and shear rate, the increase in viscosity at higher Hct outweighed the reduction in *<u>S</u>_max_*, resulting in higher *<u>τ</u>_max_*.

### Shear exposure at a given flow or oxygen delivery depends on pump operating strategy

We compared shear generation across operating conditions to determine whether CPB settings could reduce shear while maintaining flow or oxygen-delivery targets. Peak shear rate generally increased with cycle-averaged flow (Figure 4A), but differences in occlusion and hematocrit produced substantial variation in *<u>S</u>_max_* at a given *<u>Q</u>*. For example, at *<u>Q</u>* ≈1.5 cm^3^/s, *<u>S</u>_max_* ranged from approximately 1.0 to 1.7 × 10⁴ s^-1^ across the simulated operating conditions.

**Figure 4.**
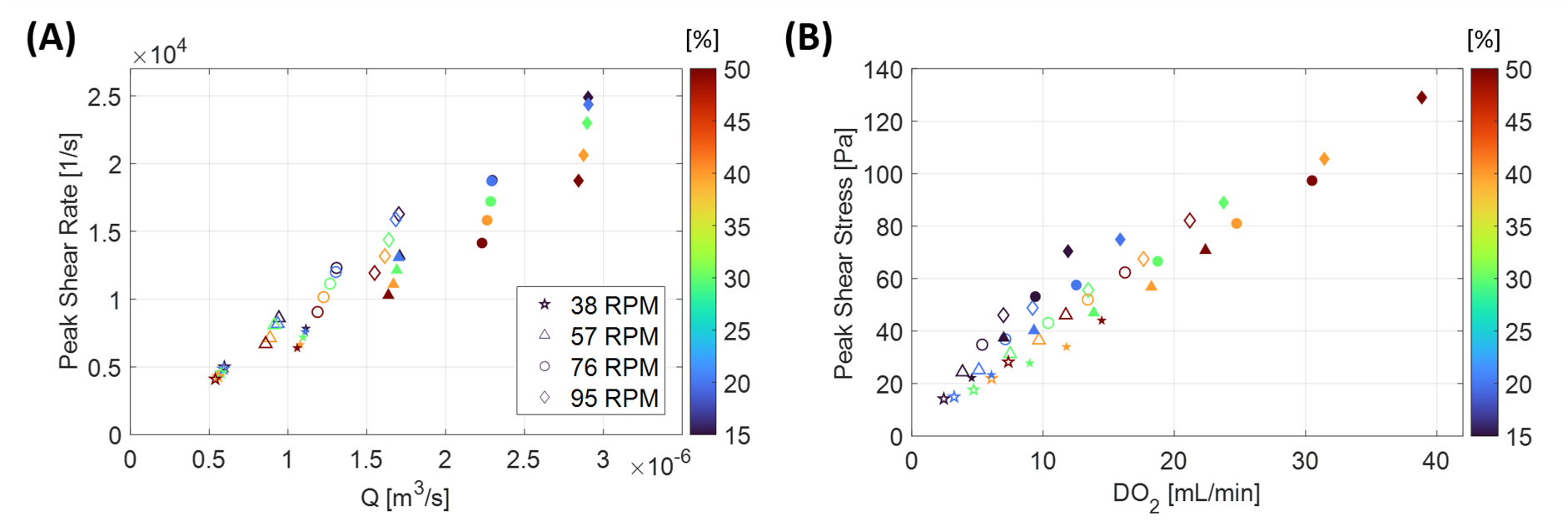
Relationships between flow rate, oxygen delivery, and flow shear across CPB operating conditions. **(A)** Cycle-averaged peak shear rate (*<u>S</u>_max_*, s⁻¹) vs. flow rate, (*<u>Q</u>*, m³/s). **(B)** Cycle-averaged peak shear stress (*<u>τ</u>_max_*, Pa) vs. oxygen delivery (DO₂, mL/min). Symbol shape denotes roller rotational speed (rpm), symbol color indicates hematocrit level, and open and filled symbols represent 77% and 83% occlusion, respectively.

Accounting for hematocrit-dependent oxygen-carrying capacity and viscosity produced a more complex relationship between oxygen delivery and peak shear stress (Figure 4B). As with flow rate, similar oxygen delivery (DO_2_) could arise from multiple operating conditions associated with markedly different peak shear stresses. For example, ten conditions yielded DO₂ values between approximately 9 and 12 mL/min, yet *<u>τ</u>_max_* ranged from approximately 25 to 70 Pa—a nearly threefold difference over a relatively narrow oxygen-delivery range. Conversely, comparable peak shear stresses occurred across a wide range of oxygen delivery: seven conditions produced *<u>τ</u>_max_* values between 70 and 80 Pa while DO₂ ranged from approximately 12 to 25 mL/min.

The reduced spread at DO₂ values above approximately 20 mL/min should not be interpreted as convergence of the oxygen delivery-peak shear stress relationship. Instead, it reflects sparse sampling of the operating space at higher oxygen delivery, which was reached by only a few parameter combinations. Additional cases with higher occlusion and/or pump speeds would likely populate this region and reveal a wider range of peak shear stresses at comparable oxygen delivery, extending the trend observed at lower DO₂.

Peak shear stress alone does not capture the unphysiological recirculation and stagnation on either side of the roller occlusion (Figure 2D). We therefore quantified these disturbances using modified relative residence time (μRRT), a metric combines wall shear magnitude and directional oscillation, that characterizes prolonged residence and poor washout near walls^43^.

Figures 5A-B show full-cycle μRRT distributions for representative high-occlusion/high-speed and low-occlusion/low-speed cases from Figure 2D. Because they span the full roller cycle, these maps describe cumulative transport, and elevated μRRT indicates longer residence inside the recirculating regions. In both cases, elevated values were concentrated within the raceway above the gray dashed line, whereas μRRT was substantially lower in the free tubing.

**Figure 5.**
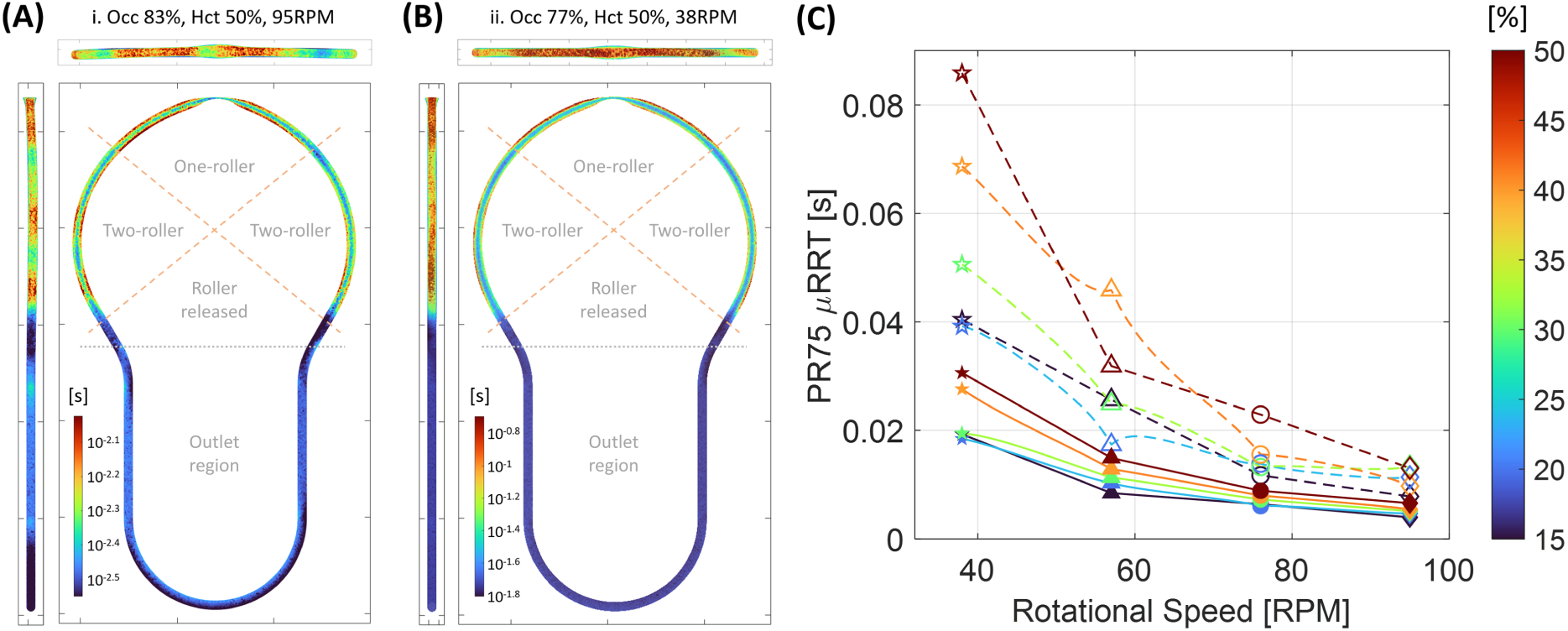
Spatial distribution and parametric dependence of relative residence time. **(A, B)** Front, top, and side view maps of μRRT (s) at the tubing wall for the same representative operating conditions shown in Figure 2D: (A) 83% occlusion, 50% hematocrit, and 95 rpm; (B) 77% occlusion, 50% hematocrit, and 38 rpm. The gray dashed line separates the raceway and free tubing segments. Within the raceway, the upper, lateral and lower quadrants correspond respectively to one-roller contact, two-roller contact, and roller release. A logarithmic colormap was used to aid visualization. **(C)** 75th percentile of μRRT within the raceway (PR75, s) vs. rotational speed (*Ω*, rpm). Symbol color denotes hematocrit; open and filled symbols represent 77% and 83% occlusion, respectively.

Within the raceway, μRRT depended on roller-contact state. Values were highest in the upper raceway, where one roller compressed the tubing during part of the cycle, and lower in the lateral raceway, where both rollers compressed it from opposite sides, although they remained above free-tubing values. Spatial patterns were similar across operating conditions, but magnitudes differed substantially. Maximum μRRT in the low-speed, low-occlusion case (*ϕ* = 77%, Hct=50%, *Ω* = 38 rpm) was approximately 2.5-fold higher than in the high-speed, high-occlusion case (*ϕ* = 83%, Hct=50%, *Ω* = 95 rpm).

Figure 5C summarizes these effects by plotting the 75th percentile of μRRT against roller speed for all occlusion-hematocrit combinations. μRRT decreased with increasing speed and occlusion, indicating improved washout. Hematocrit had a stronger effect for *ϕ* =77% than for *ϕ* = 83%, with lower values generally reducing μRRT through viscosity-dependent changes in the flow.

### Lower hematocrit permits favorable shear-residence-time trade-offs under an oxygen-delivery constraint

Because peak shear stress and relative residence time responded differently to operating conditions, we evaluated them jointly by plotting *<u>τ</u>_max_* against the 75th percentile of μRRT (PR75) across all simulations (Figure 6A). Lower PR75 μRRT generally coincided with higher *<u>τ</u>_max_*, revealing a trade-off between washout and shear. This relationship depended strongly on hematocrit: lower-Hct curves shifted toward lower *<u>τ</u>_max_* and PR75 μRRT, closer to the lower-left region where both metrics are minimized.

**Figure 6.**
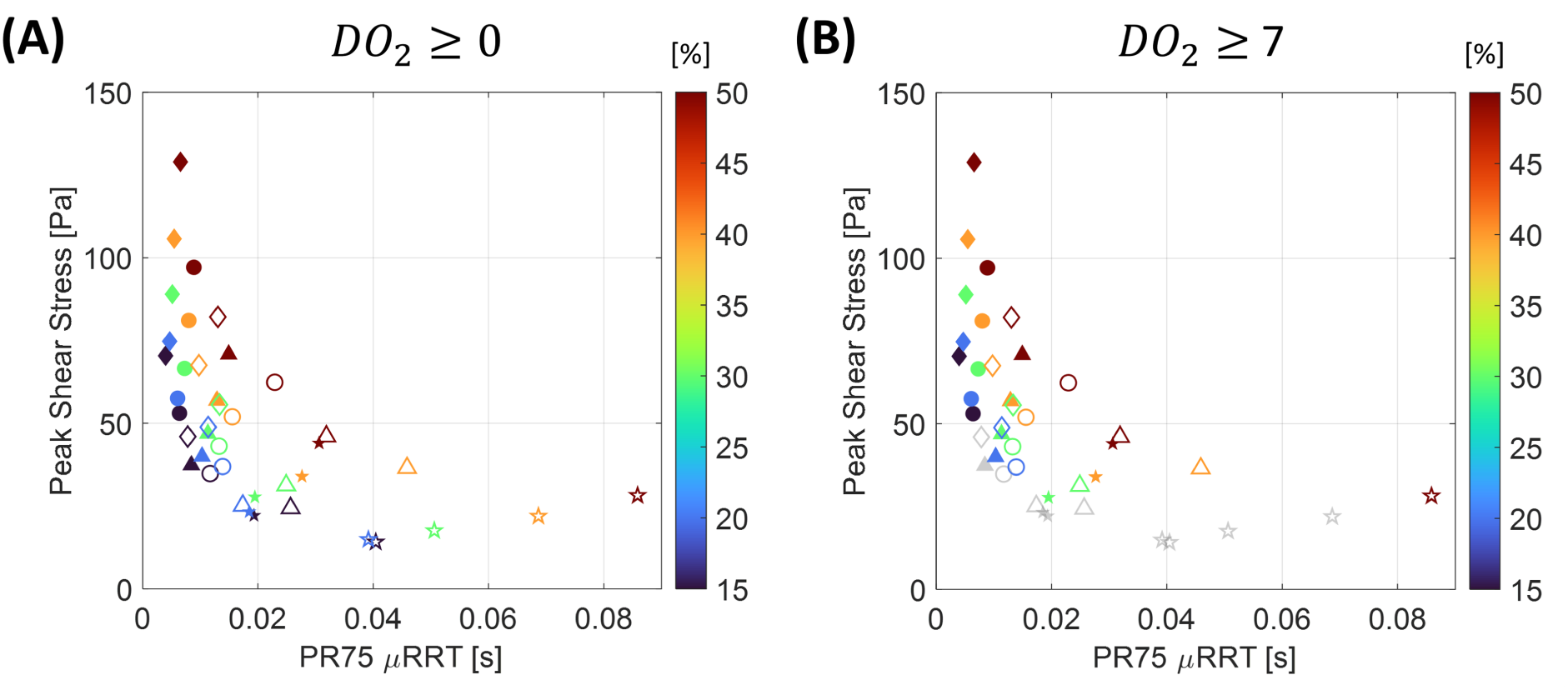
Trade-off between peak shear stress and relative residence time. **(A)** Cycle-averaged peak shear stress (*<u>τ</u>_max_*, Pa) vs. the 75th percentile of μRRT within the raceway (PR75, s). **(B)** Same plot after applying an oxygen-delivery filter (DO₂ ≥ 7 mL/min), with operating conditions below the threshold shown in gray. Symbol shape denotes roller rotational speed, symbol color denotes hematocrit level, and open and filled symbols represent 77% and 83% occlusion, respectively, consistent with previous figures.

To determine whether the favorable shift at lower hematocrit persisted after accounting for the concomitant reduction in oxygen delivery, we imposed an illustrative threshold of DO₂ ≥ 7 mL/min, selected to retain sufficient coverage of the simulated parameter space rather than to represent a clinically relevant target. The filter excluded cases with insufficient oxygen transport, shown in light gray in Figure 6B. Among the retained cases, lower-Hct conditions remained closest to the low-shear, low-residence-time region. Thus, lower hematocrit permitted occlusion and pump speed to be adjusted toward lower shear and residence time while remaining above the illustrative oxygen-delivery threshold.

## DISCUSSION

Cardiopulmonary bypass (CPB) is associated with postoperative complications, including systemic inflammatory response syndrome (SIRS) and multiorgan dysfunction^3,4^. Proposed contributors include blood contact with artificial surfaces, temperature changes, ischemia-reperfusion injury, nonpulsatile flow, transfusion, hemolysis, and supraphysiological shear stress^3–11^.

Maintaining adequate oxygen delivery (DO₂) is a central goal of CPB, yet proposed nadir indexed oxygen delivery (DO₂i) thresholds for limiting acute kidney injury vary from 258 to 400 mL/min/m² across adult and pediatric cohorts, depending on patient age and phase of bypass^18–22^. Reported hematocrit (Hct) targets are similarly heterogeneous. Two randomized infant trials compared lower and higher hematocrit strategies during hypothermic CPB^23,24^. A pooled analysis of these trials found that psychomotor scores increased with hematocrit up to 23.5% and plateaued above that level^25^. Adult studies associated thresholds of 23% and 24% with renal failure and low-output syndrome, respectively^26^. High-flow/high-Hct pediatric protocols (200 mL/kg/min, >32%) were associated with lower acute kidney injury and acute neurological event rates than conventional conditions (150 mL/kg/min, >25%)^27,28^. Conversely, Reagor et al. reported a lower nadir Hct in their high-flow than low-flow group (25% vs 28%)^29^, underscoring that flow and Hct cannot be considered independently.

This variability reflects institutional strategies that adjust hematocrit and flow primarily to preserve oxygen delivery, with limited consideration of shear in the pump circuit. Studies have shown that monocytes exposed to CPB-related shear stress upregulate inflammatory cytokines, particularly within the high-shear environment of the pump raceway^13,14,16,17^. More broadly, excessive shear has been linked to hemolysis, sublethal red blood cell damage, and leukocyte and platelet activation^44–54^. Dextran-based priming strategies have also been reported to reduce hemolysis, potentially by increasing the resistance of RBCs to shear compared to crystalloid-based priming^55^. Together, these observations expose a major gap: no established framework integrates oxygen-delivery and shear-related blood-injury criteria when selecting CPB settings.

Roller pumps remain widely used in pediatric CPB because they enable compact, low-prime circuits that limit hemodilution and transfusion exposure in infants and small children^2^. To date, comparative studies between roller and centrifugal pumps have not established clear superiority of either pump type^56–59^. Roller-pump fluid mechanics have traditionally been interpreted through classical studies of periodic peristaltic wave trains. These studies showed that forward transport is accompanied by retrograde leakage through the residual gap and fluid trapping near the moving constriction, with net flow governed by occlusion, wave speed, pressure gradient, and viscosity^30,34,35^. Asymmetric and single-wave studies, which more closely resemble roller-pump operation, confirmed these trends^60,61^. However, these idealized models generally assume viscous-dominated flow and do not resolve the detailed flow generated by realistic three-dimensional tubing deformation, for which fluid inertia may not be negligible.

Only a few computational studies have examined flow within roller pumps used for CPB. Lumped-parameter models representing the pump as a series of interconnected compartments have been used to study pressure pulsations and their dependence on pump speed, occlusion, and downstream resistance^36,37^. However, such models cannot resolve local flow structures or shear stresses within the tubing. Computational fluid dynamics models have also been reported, but these relied on two-dimensional domains reconstructed from photographed tubing profiles or prescribed using simplified geometries^31–33^. A three-dimensional FSI model was previously presented for a generic peristaltic pump^38^. In this framework, tubing deformation and the resulting flow field emerge from the coupled roller-tube-fluid dynamics rather than from a prescribed lumen geometry or inlet waveform. However, that study demonstrated feasibility rather than systematically evaluating CPB-relevant hemodynamics.

To our knowledge, this is the first pediatric CPB study to combine three-dimensional roller-tube FSI with a closed-loop circuit and systematically vary roller speed, occlusion, and hematocrit. Our three-dimensional simulations reproduced the principal flow features reported in previous two-dimensional models, including strong acceleration and elevated shear within the occluded gap, retrograde leakage beneath the roller, and substantial changes in the local flow as the rollers entered and left the raceway^31–33^. Previous studies prescribed two-dimensional compressed-lumen geometries, whereas our model captured lateral widening of the flattened tubing, partially preserving the occluded flow area and likely contributing to lower peak velocities. Our configuration also operated at lower Reynolds numbers than the previously modeled systems, consistent with the smaller tubing dimensions of pediatric CPB. Although two earlier studies used turbulence closures^32,33^, sustained turbulence at their reported Reynolds numbers under pulsatile conditions remains uncertain.

An important implication of our simulations, together with prior studies, is that roller-pump operation generates spatially and temporally heterogeneous hemodynamic insults within the raceway tubing. Blood cells passing through the occluded gap experience brief supraphysiological shear, whereas those trapped in the adjacent recirculation regions may undergo poor washout and repeated exposures to disturbed flow. Peak shear stress and relative residence time provide complementary measures of these phenomena.

A central finding of this study is that a prescribed oxygen-delivery target does not define a unique roller-pump operating condition. Roller speed, occlusion, and hematocrit affected pump performance through distinct mechanisms: roller speed primarily regulated bulk flow, whereas hematocrit had comparatively little effect on flow rate but strongly influenced oxygen-carrying capacity and shear stress through its effect on blood viscosity. This interplay creates an opportunity to optimize CPB settings beyond oxygen delivery alone. However, shear and washout cannot generally be optimized simultaneously. For example, decreasing roller speed reduced peak shear but increased relative residence time.

Among the simulated conditions satisfying the oxygen-delivery constraint, lower hematocrit permitted roller speed and occlusion to be adjusted toward a more favorable balance of peak shear stress and relative residence time. This finding is relevant to the growing use of clear crystalloid priming and transfusion-sparing strategies in pediatric CPB^62–65^. Although clear priming is intended primarily to reduce allogeneic transfusion and its inflammatory burden, it may therefore provide an additional hemodynamic benefit.

This study has limitations. Numerical constraints on roller-tube contact limited achievable residual gaps to approximately 1 mm. In the smaller pediatric tubing, this corresponded to normalized occlusions of 77%-83%—below the approximately 90% modeled previously^31–33^—and yielded flow rates below typical clinical values. Direct comparison is further limited because clinical occlusion is rarely reported despite standardized retrograde-flow setting procedures^66^. Future models should extend to higher occlusions to determine whether the observed trends persist under clinically representative flow conditions.

Peak shear stress identifies the most intense local loading, but blood-cell injury depends on both shear magnitude and exposure duration^67^. We thus complemented peak shear stress with RRT to identify regions of poor washout and prolonged residence, but RRT is not a direct measure of cumulative cell damage. More mechanistic estimates would involve accumulating an empirically calibrated damage variable along cell trajectories^68–70^.

The simplified closed-loop circuit, consistent with prior peristaltic models^71^, generated flow from roller-induced tubing deformation without prescribed inlet or outlet conditions, while omitting the full tubing length and downstream hydraulic resistance. Separately, the model neglected frictional heating, tubing fatigue, and plasticizer leaching^72,73^. Although fatigue and failure are mainly associated with prolonged pumping or excessive occlusion^74,75^, their dependence on roller speed and occlusion may impose additional optimization constraints.

## CONCLUSION

To our knowledge, this is the first systematic three-dimensional FSI study of pediatric roller-pump CPB across 40 combinations of roller speed, occlusion, and hematocrit. The simulations revealed localized high shear within the occluded gap and poor washout in adjacent recirculation regions. Different parameter combinations met similar oxygen-delivery targets while yielding distinct shear and washout environments. Lower hematocrit permitted selection of roller-speed and occlusion settings yielding lower peak shear stress and relative residence time. These findings support multimetric optimization of roller-pump operation and motivate future experimental validation under clinically representative conditions.

## Institutional Review Board Statement

Institutional review board approval was not required because this study involved computational simulations without human participants or identifiable patient data.

## Glossary of Abbreviations and Symbols

APDL: Ansys Parametric Design Language
CFD: Computational fluid dynamics
CFL: Courant-Friedrichs-Lewy
CPB: Cardiopulmonary bypass
DO_2_: Oxygen delivery
DO_2_i: Indexed oxygen delivery
FSI: Fluid-structure interaction
Hct: Hematocrit
MUSCL: Monotone Upstream-Centered Schemes for Conservation Laws
OSI: Oscillatory shear index
PISO: Pressure-Implicit with Splitting of Operators
PR75: 75^th^ percentile
PRESTO!: Pressure Staggering Option
PVC: Polyvinyl chloride
RBCs: Red blood cells
RP-CPB: Roller-pump cardiopulmonary bypass
rpm: Revolutions per minute
RRT: Relative residence time
SIRS: Systemic inflammatory response syndrome
TAWSR: Time-averaged wall shear rate
TAWSS: Time-averaged wall shear stress
*μ*RRT: Modified relative residence time
Ω: Roller rotational speed
*ϕ*: Tubing occlusion
*<u>Q</u>*: Cycle-averaged volumetric flow rate
*<u>S</u>_max_*: Cycle-averaged peak shear rate
*<u>τ</u>_max_*: Cycle-averaged peak shear stress
*μ*: Dynamic viscosity
*τ_s_*: Flow shear stress
*γ̇*: Shear rate
*D̿*: Strain-rate tensor

## Central Message

Comparable levels of oxygen delivery across pediatric roller-pump cardiopulmonary bypass settings can conceal markedly different flow shear and washout environments.

## Perspective Statement

Pediatric bypass strategies vary, and their mechanical effects remain incompletely understood. Three-dimensional fluid–structure interaction simulations link operating conditions to hemodynamics, flow shear, and washout, revealing different biomechanical environments at similar oxygen delivery. This platform enables evaluation of perfusion trade-offs to inform strategies for reducing blood trauma.

---

**Supplementary Figure 1.**
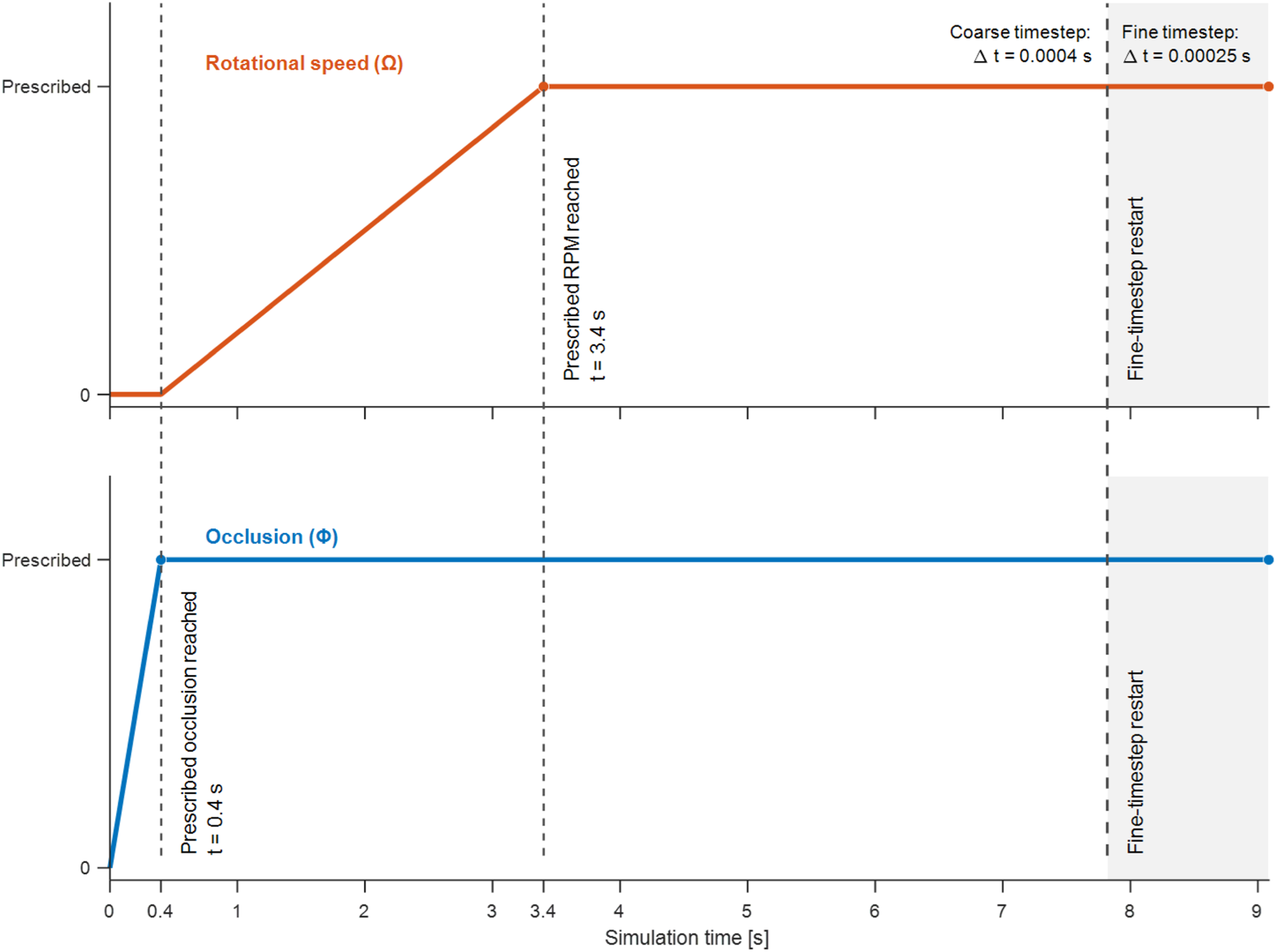
Prescribed roller motion and simulation time-stepping sequence. Initial roller engagement establishes the target tubing occlusion during 0–0.4 s. Angular speed then increases linearly from zero to the prescribed value during 0.4–3.4 s, followed by constant-speed rotation. Vertical dashed lines mark completion of initial engagement, attainment of the prescribed angular speed, and the fine-timestep restart. The shaded region indicates the fine-timestep simulation interval; the restart time varies by case. Ω, angular speed; Φ, occlusion; Δt, simulation time step.

